# Human SRP Subunits Coordinate Protein Targeting and mRNA Quality Control

**DOI:** 10.64898/2026.07.31.742061

**Authors:** Morgana K. Kellogg, Elena B. Tikhonova, Paul F. D’Cunha, Andrey L. Karamyshev

## Abstract

The Signal Recognition Particle (SRP) targets secretory proteins to the endoplasmic reticulum (ER) for their subsequent transport and protects their mRNAs from degradation by the RAPP pathway. The SRP is an RNA-protein complex consisting of six protein subunits and one noncoding RNA assembled into S- and Alu-domains. However, the distinct roles of individual SRP subunits remain undetermined. Using a molecular dissection of the human SRP through the depletion of specific subunits, we demonstrate that S-domain subunits have a primary role in mRNA protection and protein targeting, while Alu-domain subunits are dispensable. The SRP54 subunit is essential for both processes and ribosome association, while other S-domain subunits (SRP19, SRP68, and SRP72) serve supportive roles. The expression of SRP subunits is dependent on each other within the same domains, suggesting independent domain assembly. These data provide new insights into the multidomain organization and distinct functions of human SRP subunits and establish a framework for understanding the molecular mechanisms of SRP-related human diseases.

## Introduction

Protein biogenesis involves balancing the synthesis and degradation of new proteins, protein targeting, post-translational modifications, and further processing. An important aspect of protein biogenesis is correctly targeting proteins to their subcellular localization and secretion outside of the cell. These proteins comprise a secretome. The human secretome is defined as blood proteins, locally secreted proteins, and intracellular or membrane-bound proteins with at least one isoform that is secreted (Uhlen et al., 2019). It is estimated that the secretome could comprise as much as 38% of all human protein coding genes (Aebersold et al., 2018; Uhlen et al., 2015). Targeting of the secretome proteins occurs either post-translationally, after the ribosome has finished synthesizing the nascent protein, or co-translationally, when the ribosome is targeted to the endoplasmic reticulum (ER) during translation. Several post-translational mechanisms are known; they involve chaperones, ribosome-associated factors, TRC40, components of the SND pathway, and SEC62/SEC63 and others. The co-translational targeting pathway involves Signal Recognition Particle (SRP) as a major targeting factor; this pathway is known as the SRP-dependent pathway. SRP principally targets a large portion of the human secretome (Miller et al., 2026). Defects in SRP or in its substrates lead to many human diseases (Bellanne-Chantelot et al., 2018; Carapito et al., 2017; Carden et al., 2018; Goldberg et al., 2020; Gutierrez Guarnizo et al., 2023; Hernandez et al., 2021; Juaire et al., 2021; Karamyshev et al., 2020; Karamysheva et al., 2019; Kellogg et al., 2022; Miller et al., 2024; Pinarbasi et al., 2018; Schmaltz-Panneau et al., 2021; Schurch et al., 2021; Tikhonova et al., 2019).

Mammalian SRP is a ribonuclear protein complex consisting of six protein subunits and one structural, long, noncoding RNA called 7SL (Kellogg et al., 2021), shown in **Fig. 1A**. The SRP protein subunits are categorized into two domains that occupy about half of 7SL RNA each: the Alu domain (heterodimer SRP9 and SRP14), which is involved in translational control (Lakkaraju et al., 2008; Mary et al., 2010; Siegel and Walter, 1986), and the signal recognition or S-domain (SRP19, SRP54, and a heterodimer of SRP68 and SRP72) (Brown et al., 1994; Lutcke et al., 1992; Siegel and Walter, 1988).

**Fig. 1.**
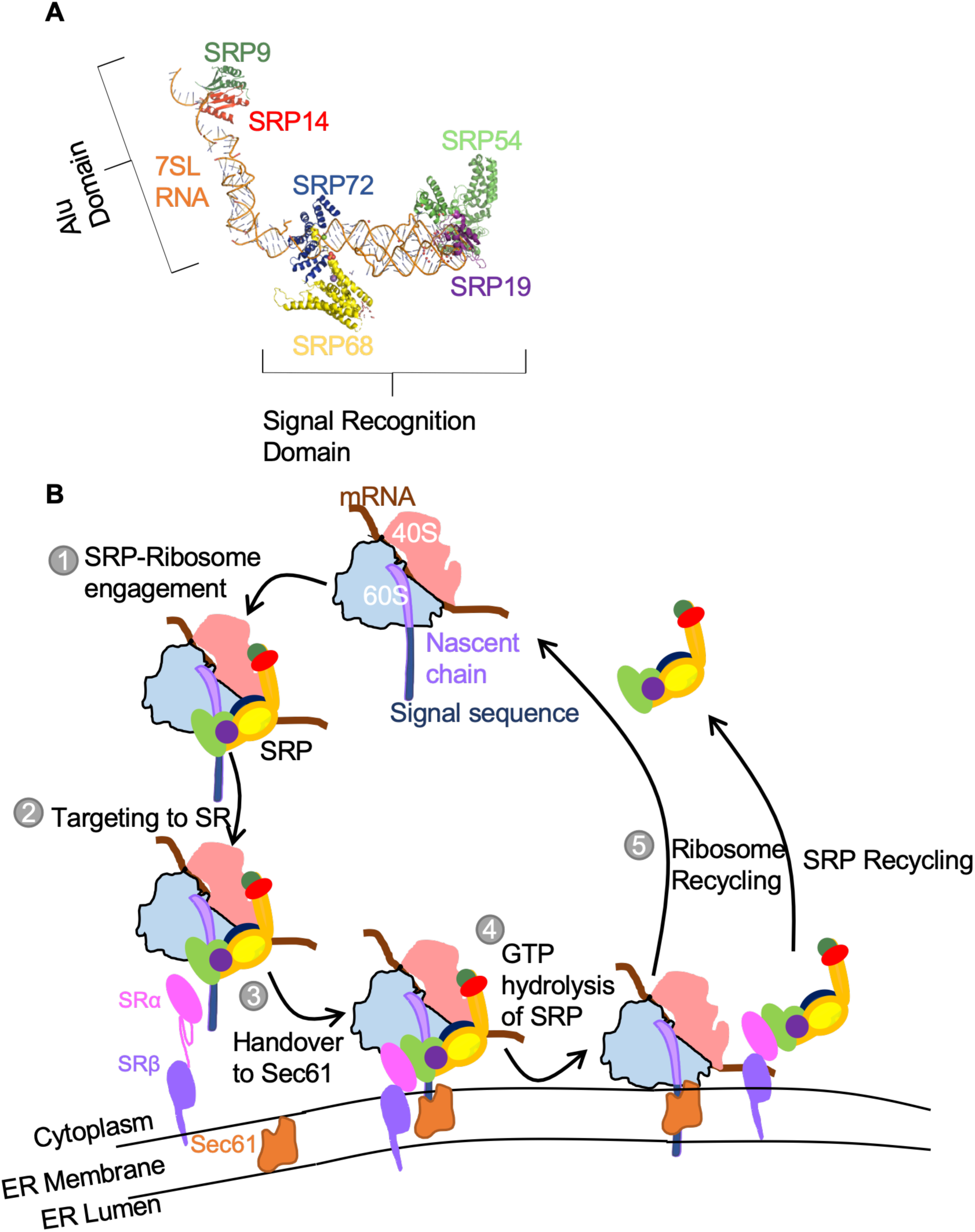
SRP and the SRP Cycle. **A**, SRP is a six-protein subunit complex arranged on 7SL noncoding RNA. There are six protein subunits named by their molecular weights: SRP9 and SRP14 in the Alu domain (green and red respectively); SRP19 (purple), SRP68 (yellow), SRP72 (dark blue), and SRP54 (light green) in the signal recognition or S domain. The SRP model was based upon (Kellogg et al., 2021). **B**, Schematic Representation of the SRP Cycle. In SRP-mediated protein targeting, the SRP recognizes an N-terminal signal sequence of the nascent polypeptide chain (light purple) called the signal sequence (navy extension). The SRP engages the ribosome (light pink and light blue structures, Step 1) and then transports the ribosome to the SRP receptor (dark pink and dark purple) on the ER lumen (Step 2), where the ribosome is handed over by the receptor to the Sec61 translocon (orange) to continue translation (Step 3). SRP is then recycled to produce and target other proteins (Steps 4-5 correspondingly). All SRP subunit colors are the same as in panel A.

How SRP biogenesis occurs in mammals is not completely understood. Most available information infers function from different model systems, including yeast, *Plasmodium falciparum*, and *Escherichia coli* (Brooks et al., 2009; Panchal et al., 2014; Poritz et al., 1990; Romisch et al., 1989; Soni et al., 2021; Tuteja, 2007). Others have measured the binding between mammalian SRP subunits using purified human protein complex *in vitro* or have studied the structure of zebrafish, mammalian, or yeast SRP (Becker et al., 2017; Brooks et al., 2009; Carapito et al., 2017; Egea et al., 2008; Grotwinkel et al., 2014; Hainzl et al., 2002; Hainzl et al., 2007; Iakhiaeva et al., 2009; Juaire et al., 2021; Kobayashi et al., 2018; Oubridge et al., 2002; Schurch et al., 2021; Wild et al., 2016; Wild et al., 2019). Few studies have examined SRP subunits endogenously in a human cellular system; Juaire et al tested SRP54 clinical mutations in HEK293 cells (Juaire et al., 2021); and Linder et al. studied the granulopoiesis of neutrophils in SRP19 genetic defects in inducible human pluripotent hematopoietic stem cells (Linder et al., 2023).

The most commonly accepted *in vitro* model of SRP biogenesis has several steps: first, 7SL RNA is transcribed in the nucleus by RNA polymerase III (Elder et al., 1981), while SRP protein subunits are synthesized in the cytoplasm. Next, all SRP subunits except SRP54 are imported into the nucleus and targeted to the nucleolus. SRP19 then binds 7SL to promote the binding of the heterodimers SRP68/72 and SRP9/14. These five SRP subunits with 7SL form a pre-SRP complex, which is then exported from the nucleus. Finally, SRP54 is loaded to the pre- SRP particle with assistance of SMN (Survival of Motor Neuron) complex finalizing the SRP maturation step (Issa et al., 2024; Piazzon et al., 2013). More details about *in vitro* SRP structure and biogenesis can be found in the review (Kellogg et al., 2021).

Once SRP has been assembled, it can then target proteins with signal peptides and transmembrane domains via the co-translational protein targeting process, known as the SRP cycle (**Fig. 1B**). Briefly, there are five general steps to the SRP cycle: recognition of the signal sequence or transmembrane domain (TMD) as it emerges from the ribosome tunnel during its synthesis and SRP association with ribosomes (**Fig. 1B** Step 1); targeting of the SRP-ribosome-nascent chain complex (SRP-RNC) to the SRP receptor (SR) on the endoplasmic reticulum membrane (**Fig. 1B** Step 2); handover of the SRP-RNC to the translocon (**Fig. 1B** Step 3); GTP hydrolysis, that is needed to release SRP from the receptor on the ER and the ribosome (**Fig. 1B** Step 4); resuming of protein synthesis with translocation of the nascent peptide into the ER lumen through SEC61 translocon followed by recycling of the ribosomal subunits and SRP in the cytosol (**Fig. 1B** Step 5). Details about this process are reviewed in (Kellogg et al., 2021).

However, when SRP fails to recognize the signal sequence, either due to a mutation in the signal sequence or a defect in SRP, a quality control mechanism called the Regulation of Aberrant Protein Production (RAPP) is induced (Karamyshev and Karamysheva, 2018; Karamyshev et al., 2020; Karamysheva and Karamyshev, 2023). RAPP leads to the specific degradation of the SRP substrate mRNA with an erroneous signal sequence (Karamyshev et al., 2014; Miller et al., 2024; Pinarbasi et al., 2018; Tikhonova et al., 2019). RAPP also occurs with the loss of the SRP54 subunit itself (Karamyshev et al., 2014; Miller et al., 2026; Pinarbasi et al., 2018; Tikhonova et al., 2022). However, it is unknown whether other SRP subunits, particularly in the S-domain, influence RAPP and the ability of SRP to protect mRNA.

Complex organization of SRP and its biogenesis, its crucial function in protein transport, SRP’s new role in mRNA protection, and its association with many diseases, make SRP very attractive for studies. However, despite numerous studies, many aspects of the molecular mechanisms of the above processes remain unknown. In our study, we hypothesize that human SRP subunits have additional functions, including attenuating other subunit protein expressions, influencing ribosome association of the whole complex, and protecting SRP-dependent substrates’ mRNA. This work is the first systematic and comprehensive study of the SRP subunits’ expression, function, and their role in mRNA protection and ribosome association in human cells. Our data demonstrate that the lack of S-domain SRP subunits (SRP19, SRP68, and SRP72) attenuates SRP54 protein expression; and the integrity of the SRP9/14 and SRP68/72 dimers depends on the presence of each subunit in the complex. We found that the depletion of S-domain SRP subunits induces a loss of mRNA expression of SRP-dependent substrates.

SRP54 depletion leads to the loss of other SRP subunits co-fractionation with the ribosome. We conclude that SRP54 is essential for SRP association with the ribosome and RAPP. We further conclude that SRP19, SRP68, and SRP72 lead to SRP biogenesis defects and are all critical to the RAPP pathway and SRP-dependent protein secretion. In our studies, we used a molecular dissection of SRP by a depletion of every subunit in cultured human cells and studying their role in all above processes. This approach may help to understand molecular mechanisms of human diseases associated with defects in SRP.

## Results

### SRP subunits affect each other’s expression in S and Alu domains suggesting their independent assembly

The mammalian SRP consists of six protein subunits assembled on 7SL RNA and arranged in Alu (SRP9 and SRP14) and S (SRP19, SRP54, SRP68 and SRP72) domains **(Fig. 1)**. The organization and the assembly of this hetero-subunit complex formation is hierarchical and well regulated. However, does the subunits’ abundance depend on each other? Will the depletion of one influence the expression of the others? While these are very interesting questions to elucidate information about hetero-complexes’ formation, they are also important queries for understanding molecular mechanisms of multiple pathologies associated with SRP defects. To answer these questions, we independently depleted each SRP subunit by siRNA in cultured human HeLa Tet-On cells and analyzed all subunits by Western blotting with subsequent quantification (**Fig. 2**). Strikingly, the SRP subunits indeed depend on each other, especially when the subunits were depleted in the same domain. Depletion of any S domain subunit (SRP19, SRP68, and SRP72) leads to significant reduction of the SRP54 protein (**Fig. 2**).

**Fig. 2.**
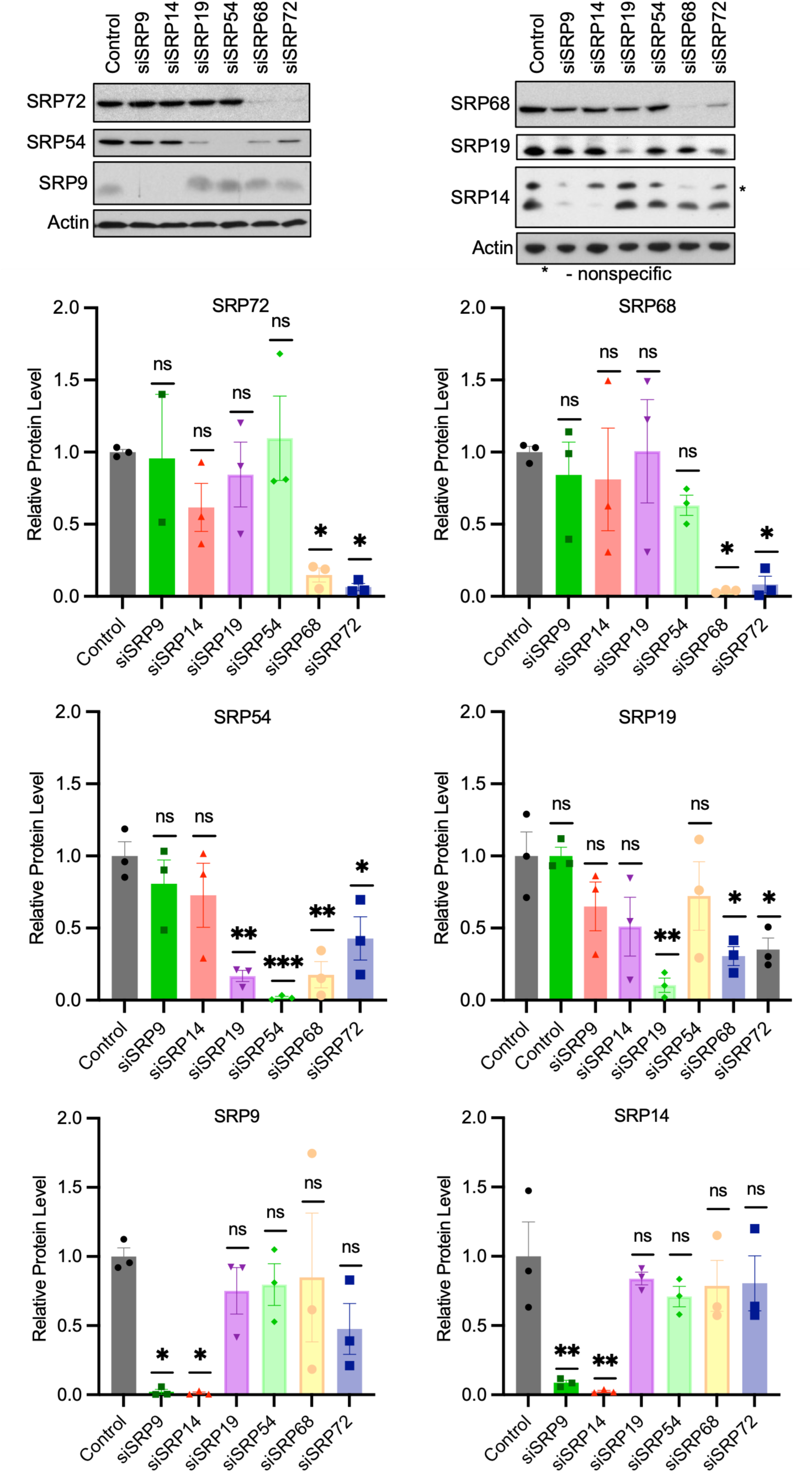
The protein expression of SRP subunits is dependent on each other within the same functional domains. Protein levels of SRP subunits were analyzed by Western blot in whole-cell lysates generated 48 hours after the corresponding siRNA transfection. Representative blots are shown. Actin was used as a loading control. Graphs show quantitative analysis based on three biological replicates and represent the mean ± the standard error. Statistical significance was determined by one-way ANOVA with post-hoc Dunnett’s test; p > 0.05, * p < 0.05, ** p<0.01, *** p<0.001, ns is not significant.

However, knockdown of SRP54 did not reduce either SRP19, SRP68 or SRP72. Depletion of SRP68 or SRP72 reduced SRP19; and the abundance of SRP68 and SRP72 strictly depends on each other. Similarly, SRP9 and SRP14 also strictly depend on each other (**Fig. 2**). SRP9/SRP14 and SRP68/SRP72 form heterodimers in the SRP structure (Birse et al., 1997; Weichenrieder et al., 2000), and our data affirm their relationship. Notably, depletion of subunits in one domain do not significantly affect subunits of another domain, suggesting that the domains assembled independently from each other.

### Negative effect of an SRP subunit depletion on other subunits is not caused by mRNA abundance

Disbalance in the complex subunits associated with depletion of one of them may be caused by changes in mRNA transcription or mRNA and protein stability. For instance, subunits of the FACT chromatin remodeling complex are regulated on both protein and mRNA levels (Safina et al., 2013). To test if depletion of an SRP subunit affects mRNA abundance of other SRP subunits, we used reverse transcription quantitative real time PCR (RT-qPCR). In these experiments, each SRP subunit was independently depleted by siRNA in the cultured human HeLa Tet-On cells. The total RNA was purified, and mRNA levels were determined for all SRP subunits (**Fig. 3**). While we have obtained a very good depletion by the corresponding siRNAs as demonstrated by measurements of their mRNAs and proteins, we have not observed any statistically significant changes in abundance of mRNAs of other SRP subunits. Thus, observed changes in protein SRP subunits are not caused by their mRNAs.

**Fig. 3.**
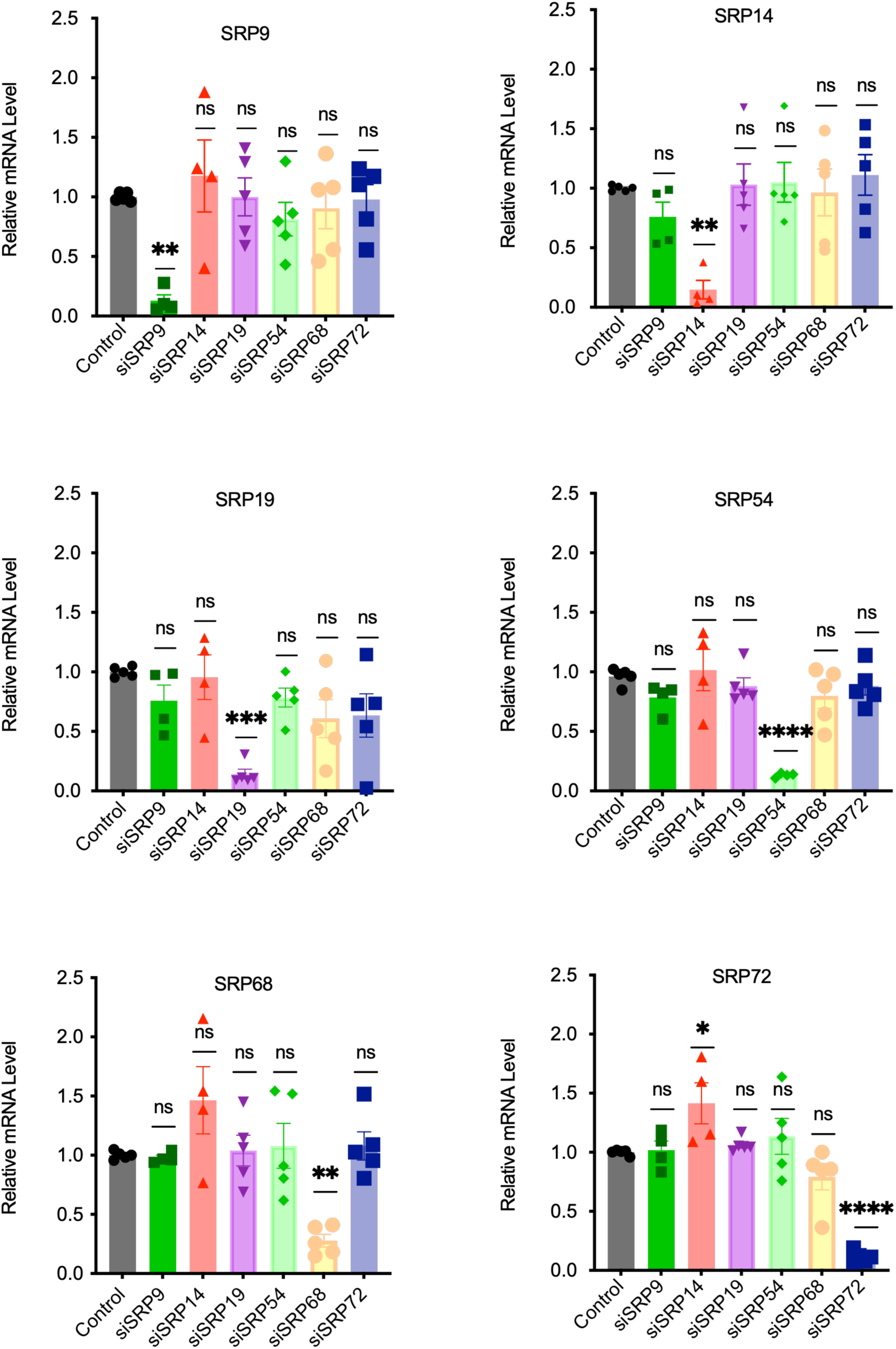
SRP subunits are not transcriptionally dependent on other subunits. HeLa Tet-On cells were cultivated for 48 hours after the corresponding siRNA transfection, total RNA was purified, cDNA synthesized and mRNA of each SRP subunit was measured by RT-qPCR. Graphs show relative levels of mRNAs calculated by comparative C_T_ method (see Material and Methods for details). Statistical significance was determined by one-way ANOVA with post-hoc Dunnett’s test, * p < 0.05, ** p<0.01, *** P<0.001, **** p<0.0001, ns is not significant. Number of replicates are indicated by individual points on each bar. The bars are the mean ± the standard error.

### Effect of inhibition of the proteasome on SRP subunits

As shown in **Fig. 2**, the S-domain proteins SRP19 and SRP54 are affected by depletion of other SRP subunits. One possible mechanism of the proteins’ reduction is degradation by the Ubiquitin-Proteasome System (Pohl and Dikic, 2019). Proteasome degrades proteins labeled by ubiquitin (especially K48-ubiquitin linked proteins) (Rieser et al., 2013). To assess whether the proteasome regulates SRP protein levels when other subunits are absent, we completed experiment with MG-132, a proteasome inhibitor, in the cultured HeLa Tet-On cells. Each SRP subunit was independently depleted using siRNA, the cells were treated with either MG-132 or DMSO, and K48-linked ubiquitination and SRP subunits were analyzed by Western blot (**Fig. 4**). As expected, MG-132 treatment leads to an increase in K48-linked ubiquitination demonstrating that proteasome degrades proteins in the cells. However, MG-132 did not affect SRP subunits - we observed no significant rescue in the heterodimer pairs (9/14 and 68/72) or SRP54 or SRP19 protein abundance (**Fig. 4**). Therefore, the protein level difference in SRP when its composition was affected was not due to degradation facilitated by the proteasome.

**Fig. 4.**
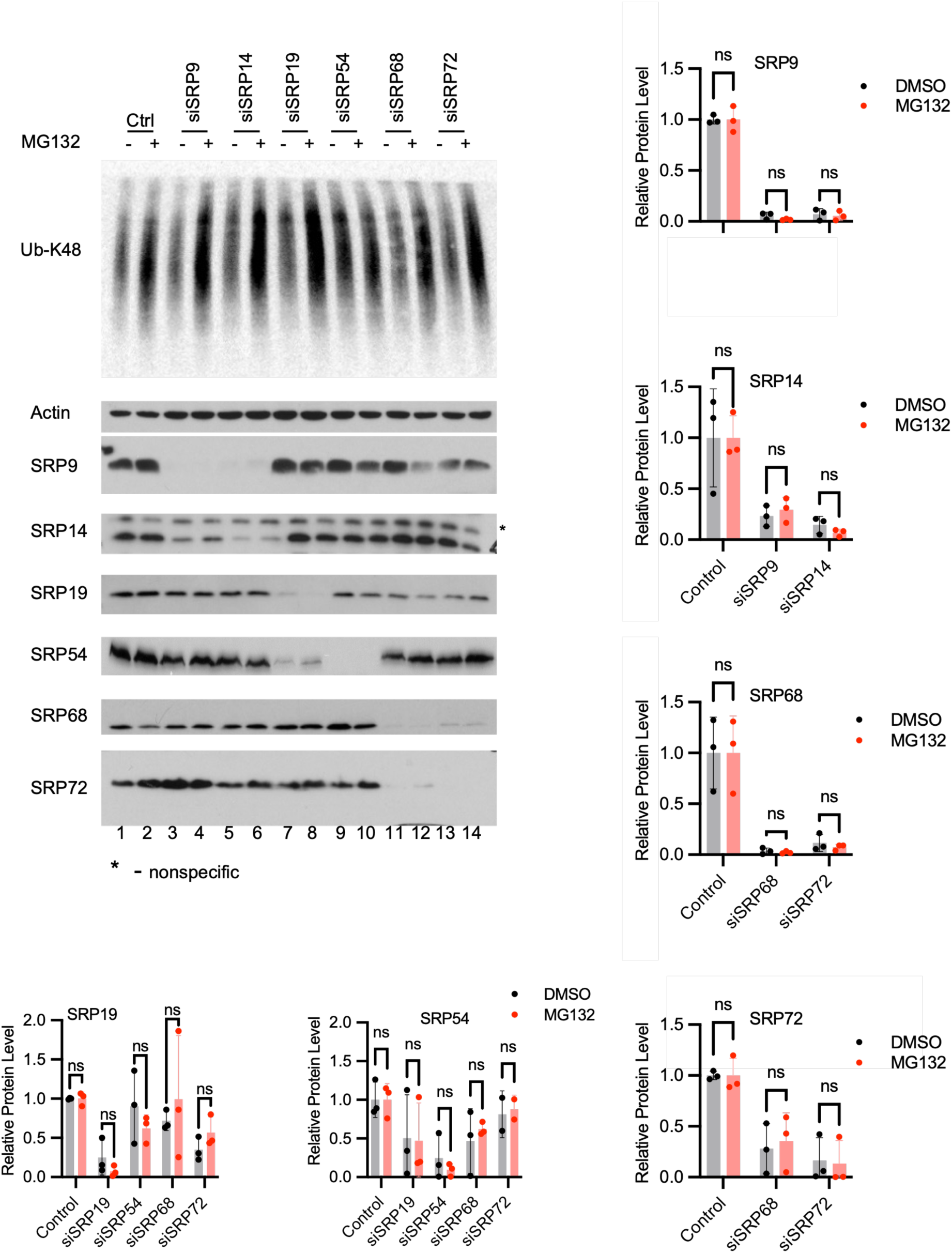
Decrease in the abundance of SRP subunits is not caused by proteasome-mediated degradation. HeLa Tet-On cells were grown, transfected with corresponding siRNAs, and treated with proteasome inhibitor MG-132 or DMSO) for approximately eight hours. The samples were analyzed by Western blot with following quantification. The representative blots are shown. K48-linked ubiquitin was used as a control for proteasome inhibition by MG132. Actin was used as a loading control. Graphs represent the mean ± the standard error on the base of three replicates. Statistical significance was determined by two-way ANOVA with post-hoc Tukey’s test for multiple comparisons; ns, not significant.

### SRP54 deficiency affects SRP association with the ribosome

We hypothesized that SRP54 depletion can negatively affect SRP association with ribosomes. To test this hypothesis, we used two alternative approaches - a crude fractionation method and polysome profiling. In the first method, clarified cellular lysates from control and SRP54 depleted cells were fractionated into two major fractions, with ribosomes precipitated by ultracentrifugation though a sucrose cushion (ribosome pellet), and a ribosome-free cytosol (supernatant after ribosome removal). As it is shown in **Fig. 5A**, this fractionation effectively separates ribosomes from the cytosol – the small subunit ribosomal protein RPS6 and the large subunit ribosomal protein RPL11 (used for ribosome detection) were found by Western blot in the pellet only, while β-actin, a cytosolic protein, was detected mostly in the supernatant. SRP54 knockdown was efficient – it was not detected by Western blot in the fractions from SRP54 depleted cells. SRP54, SRP72, and SRP14 were mostly found in the pellet, and SRP19 in the pellet and supernatant of the control cells (**Fig. 5A**). However, SRP54 depletion leads to redistribution of the SRP subunits - less SRP14, SRP19, and SRP72 proteins were associated with the pellet, and more found in the supernatant (**Fig. 5A**). These results indicated that SRP subunit interactions with the ribosome are disrupted when SRP54 is depleted.

**Fig. 5.**
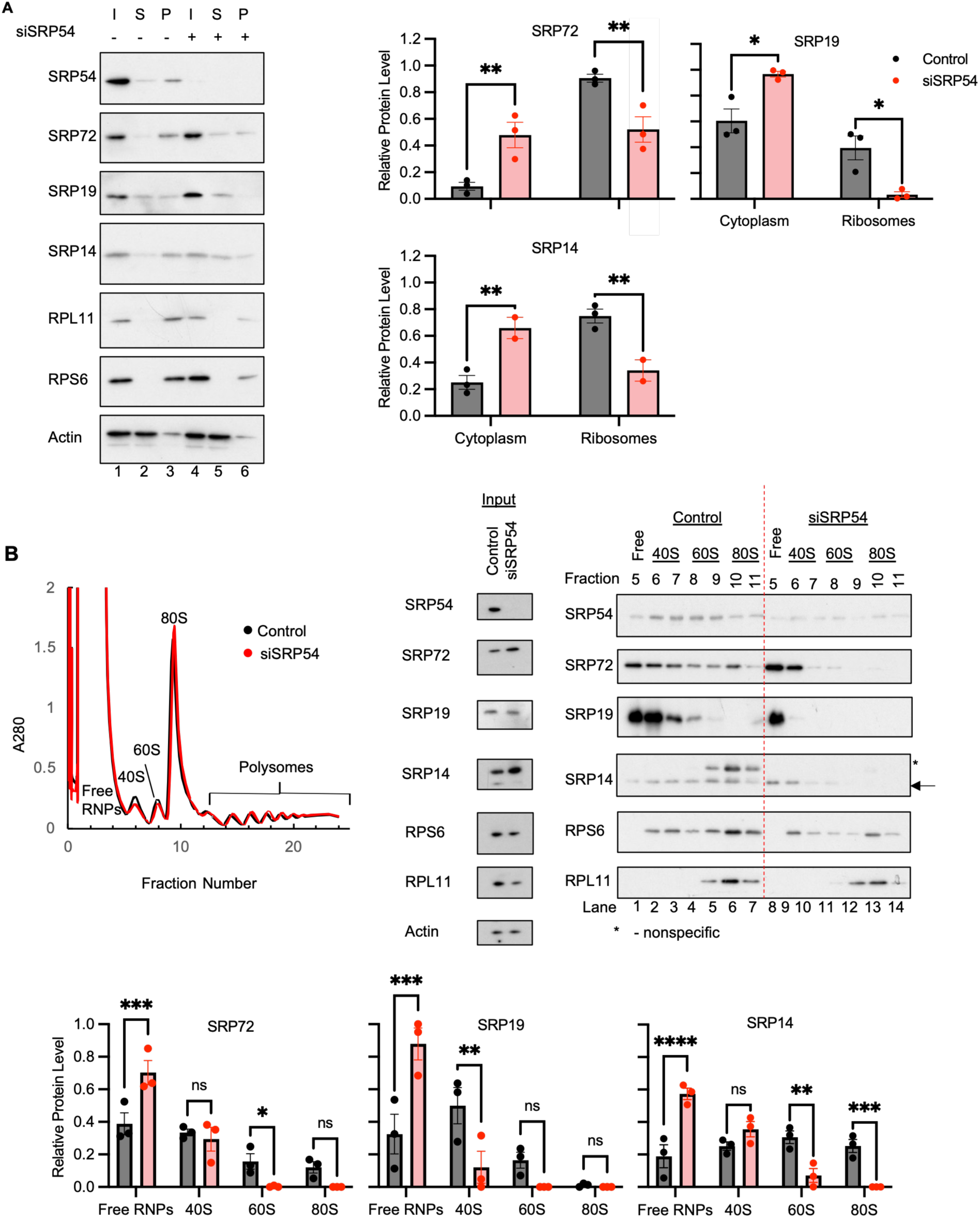
SRP54 depletion diminishes the SRP subunits association with ribosome. The association of the SRP subunits with ribosomes was tested by crude fractionation of cell lysates (**A**) or by polysome profiling (**B**) followed by Western blot analysis. SRP54 was depleted by siRNA for SRP54 and cells were treated with cycloheximide before lysis. **A**, Crude fractionation of cell lysates. SRP subunits, ribosomal proteins RPL11 and RPS6 (markers for large and small ribosome subunits, respectively), and actin (as a cytoplasmic marker) were analyzed by Western blot. Representative blots are shown. Cytoplasm and ribosome enriched fractions were separated by high-speed centrifugation on a sucrose cushion. Abbreviations: I, input (lysate before fractionation); S, supernatant (cytoplasm); P, pellet (ribosomes). siSRP treated cells are marked by “+”, controls cells by “-“. Graphs show quantitative analysis based on three biological replicates and represent the mean ± the standard error. Protein levels in the cytoplasmic and ribosome fractions were expressed as a proportion of the total level (cytoplasm plus ribosome). Statistical significance was determined a two-way ANOVA with post-hoc Dunnett’s test; ns p > 0.05, * p < 0.05, ** p < 0.01. **B**, Polysome profiling. Ribosomes and polysomes were fractionated as described in (Karamysheva et al., 2018). The representative absorbance spectra of the fractions from the siSRP54 treated cells and control are shown. Fractions corresponding to free ribonuclear proteins (free RNPs), small (40S) and large (60S) ribosome subunits, monosomes (80S) and polysomes are marked. The fractions were analyzed by Western blot similar to the panel A. Protein levels in the prepolysome fractions were expressed as a proportion of the total level (free + 40S + 60S + 80S). Statistical significance was determined by two-way ANOVA with post-hoc Dunnett’s test to compare control versus siSRP54 in GraphPad Prism; p > 0.05, * p < 0.05, ** p < 0.01, *** p < 0.001, **** p <0.0001, ns, not significant.

To independently verify these findings, we used polysome profiling (Karamysheva et al., 2018) for HeLa cells with SRP54 knockdown and control cells. This technique fractionates the cellular ribosomes into 40S and 60S ribosome subunits, 80S monosome, and polysomes (**Fig. 5B**) by ultracentrifugation in a 10-50% sucrose gradient. Similar to the above experiment, we were able to obtain very efficient depletion of SRP54 (**Fig. 5B**, Input). We also tested actin, RPS6, RPL11, SRP14, SRP19, and SRP72 in the cellular lysates before fractionation (Input), and found that they were not notably changed by SRP54 knockdown. Analysis of fractions after gradient centrifugation, in general, confirmed that SRP14, SRP19, and SRP72 subunits are redistributed from ribosome associated fractions towards fractions containing ribosome-free ribonuclear proteins (RNPs) (**Fig. 5B**). These experiments suggest that in the absence of SRP54, other SRP subunits do not associate with the 80S ribosome. We propose that SRP54 is the primary determinant for SRP-ribosome association of the whole complex in human cells.

### Integrity of the SRP S domain is required for mRNA protection of the SRP substrates

As we demonstrated earlier, in addition to its targeting function, SRP has a mRNA protection role in the cells; the loss of SRP54 triggers a specific mRNA degradation of secretory and membrane proteins in a process termed RAPP (Karamyshev et al., 2014; Pinarbasi et al., 2018; Tikhonova et al., 2022; Tikhonova et al., 2019). Do other SRP subunits (SRP9/14, SRP19, or SRP68/72) contribute to mRNA protection? What happens to the mRNAs of secretory and membrane proteins when these subunits are depleted? Previously, we completed deep RNAseq analysis of mRNAs affected by the SRP54 knockdown and found that mRNAs of many secreted and membrane proteins were depleted (Tikhonova et al., 2022). To study effects of the other SRP subunits, we selected mRNAs from these group coding proteins with different subcellular localizations: secreted proteins destined for the extracellular matrix (LGALS3BP, LOX, MMP2, SPOCK1; red bars in **Fig. 6**); endoplasmic reticulum lumen proteins (CALR, HSPA5, HYOU1, PDIA3; green bars in **Fig. 6**); plasma membrane-bound proteins (ALPI, CD24, CD248, CHRND, HLAB, KCNQ1, SLC39A14; purple bars in **Fig. 6**); and cytoplasmic or SRP-independent proteins as negative controls (HPRT, SRα, GAPDH; orange bars in **Fig. 6**). The functions of these proteins are presented in **Supplementary Tables 4 and 5**.

**Fig. 6.**
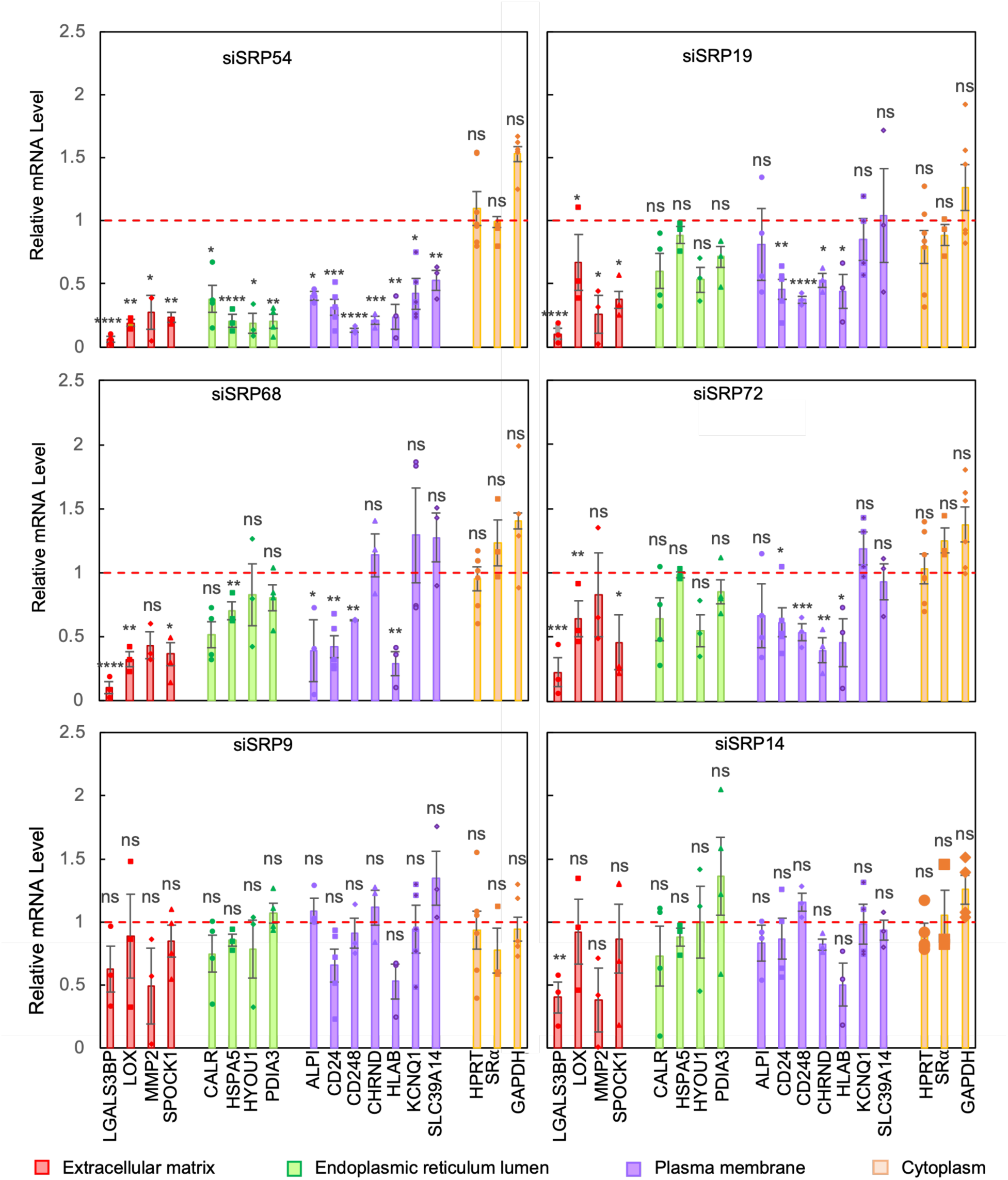
SRP S domain subunits protect specific protein mRNAs from degradation. Human HeLa Tet-On cells were cultured and transfected with siRNA against each SRP subunit. Total RNA samples were collected 72 hours after siRNA treatment of SRP. The mRNA of each subunit depletion was then quantified using RT-qPCR relative to β-Actin levels and then compared to the mRNA level of control cells. The red dashed line indicates the mRNA level in control samples. The bar graph shows mean values with standard error of the mean (SEM), number of independent biological replicates are indicated by the individual scatter points. Proteins from different cellular compartments were used for analysis. Subcellular localizations of proteins are indicated as extracellular matrix (secreted, red bars), ER lumen (green bars), plasma membrane (purple bars), and cytoplasm (negative control, orange bars). Statistical significance was determined per protein with a one-way analysis of variance (ANOVA) coupled with a post-hoc Dunnett’s test using GraphPad, * p < 0.05, ** p < 0.01, *** p <0.001, **** p < 0.0001, ns, not significant.

The RT-qPCR results show that SRP54 knockdown has the most detrimental effect of all subunits on mRNA of SRP-dependent proteins (**Fig. 6**). All of those mRNAs, except for negative controls, were significantly decreased, regardless of their final subcellular localization. Noticeable but less pronounced effects were observed for the same mRNAs when SRP19, SRP68 and SRP72 were depleted (**Fig. 6**). However, knockdowns of SRP9 or SRP14 do not significantly affect these mRNAs. The only mRNA that was affected by SRP14 knockdown was LGALS3BP. Please note, this mRNA was the most sensitive among all others to depletion of other SRP subunits. LGALS3BP has a shorter mRNA half-life (20 hours) than CALR (34 hours) and PDIA3 (26 hours) (Schwanhausser et al., 2011), so LGALS3BP mRNA may be less stable in general, and thus more sensitive to SRP subunits’ depletion.

To further probe the function of SRP subunits in protein secretion we selected representative proteins for all four classes described above: galectin-3-binding protein (LGALS3BP), which is secreted to the extracellular matrix; PDIA3, an ER lumen disulfide isomerase; SLC39A14, a divalent zinc ion symporter; CHRND, an acetylcholine receptor subunit, and beta-actin, and analyzed them in cells and medium when SRP subunits were depleted. In general, protein expression (**Fig. 7**) correlates with mRNA expression (**Fig. 6**).

**Fig. 7.**
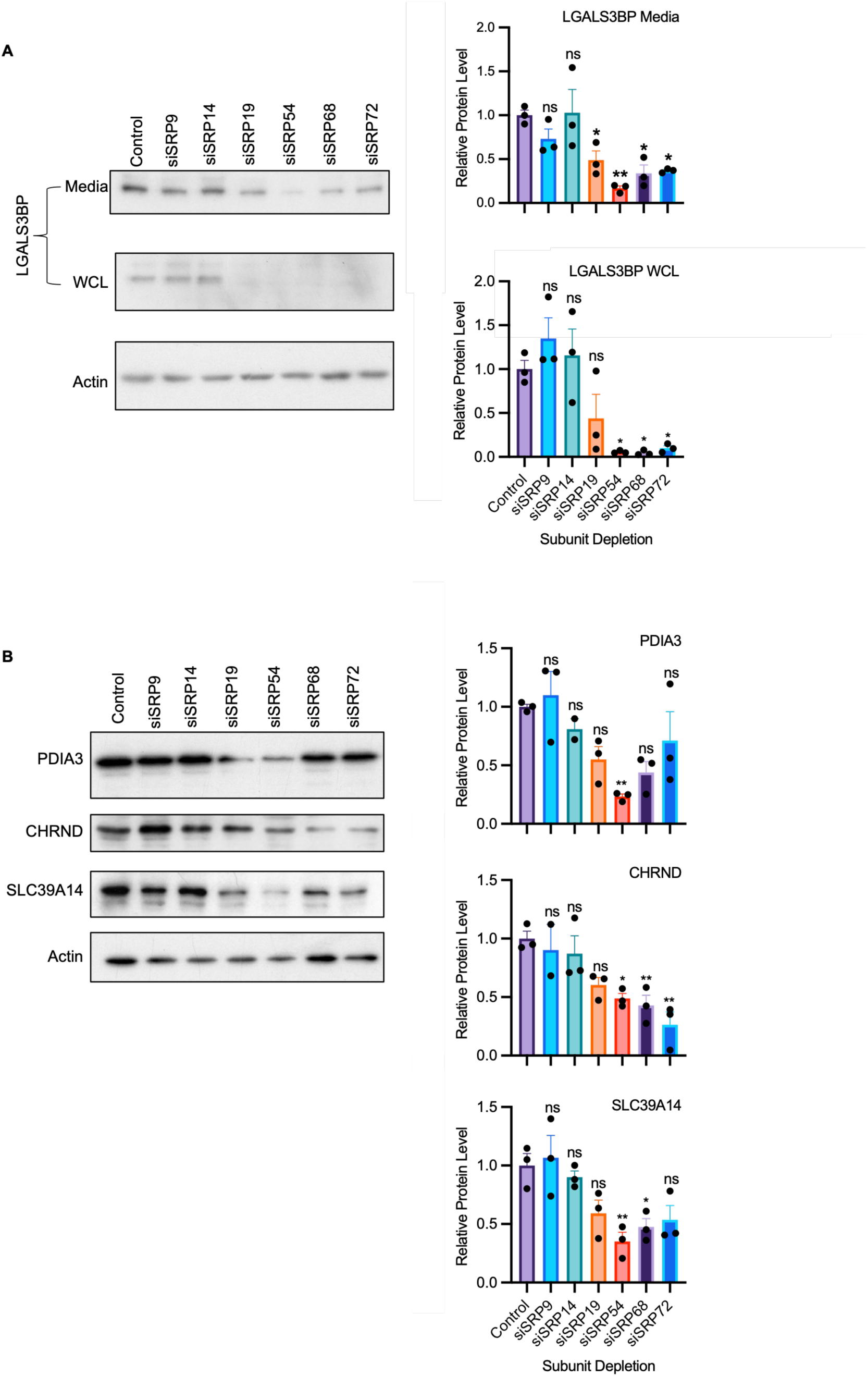
Depletion of SRP S domain subunits diminishes expression of secretory, ER and membrane proteins. Extracellular LGALS3BP protein (**A**), ER lumen protein PDIA3, and membrane proteins SLC39A14 and CHRND (**B**) were analyzed by Western blotting, followed by quantitation. SRP subunits were depleted in HeLa Tet-On cells by siRNAs. Whole cell lysate (WCL) and media (for LGALS3BP only) were collected 72 hours after siRNA transfection and analyzed by Western blotting. Representative Western blots are shown. Graphs show relative protein levels of the proteins calculated from the blots, actin was used for normalization in the lysates. The bar graphs show mean values with standard error of the mean; number of independent biological replicates are indicated by the individual scatter points. Statistical significance was determined with a one-way analysis of variance (ANOVA) coupled with a post-hoc Dunnett’s test using GraphPad, * p < 0.05, ** p < 0.01, ns, not significant.

Our results demonstrate that integrity of the S-domain, consisting of SRP54, SRP19, SRP68, and SRP72, is essential for protection of the mRNAs of membrane and secreted proteins. While SRP54 has the primary role in this process, the more remote subunits (SRP19, SRP68, and SRP72) from the ribosome exit tunnel are also significant although the effect of their depletion is lower. Most likely, the mRNA protection depends on signal peptide recognition which is regulated by the S domain of SRP. Contrary to the S domain, the Alu domain (SRP9/SRP14) is not involved in mRNA protection. It functions in translational control, not signal recognition (Bousset et al., 2014; Halic et al., 2004; Lakkaraju et al., 2008; Mary et al., 2010). The proteins of the Alu domain (SRP9/SRP14) are located a significant distance away from the signal recognition domain.

## Discussion

In this study, we examined SRP biogenesis and function by dissecting each subunit of this complex, performing detailed analysis of each subunit protein and mRNA levels, their interactions with ribosomes, their role in protein targeting, and in protecting mRNA of SRP-dependent proteins using cultured human cells as a model system mimicking *in vivo* events. It is the most comprehensive study of SRP in human cells up to date. Eukaryotic SRP consists of two major domains, S-domain and Alu-domain. It is striking that abundance of the SRP subunits in the domains depends on each other inside of the domain, but not between the domains (**Fig. 2**). Our data suggest that two distinct SRP domains assemble and function independently from each other. Indeed, S-domain subunits are the most sensitive to depletion of the others in the same domain - especially more distal subunits (SRP54, SRP19). SRP54 is affected by all S-domain subunits, and SRP19 by the depletion of SRP68 and SRP72; in contrast, the SRP central subunits, comprising the heterodimer of SRP68 and SRP72, strictly depend on each other’s expression but not on other subunits. Similarly, the Alu-domain subunits, heterodimer SRP9 and SRP14, strictly depend on each other’s abundance but are not affected by depletion of the S- domain subunits in agreement with the early observations (Gussakovsky et al., 2023). SRP54 was the most affected protein by the depletion of the other subunits of the S-domain likely because it requires the already assembled pre-SRP complex. Our findings provide new aspects of the SRP biogenesis, which has mostly been studied *in vitro* or in yeast as reviewed in (Kellogg et al., 2021).

It seems that the effects of the SRP subunits on each other’s abundance are not regulated at the mRNA level – at least we have not observed significant differences. Thus, the expression of the SRP subunits is regulated at the protein level – it could be that proteins are not stable and degrade when others are missing; alternatively, their synthesis may be affected at the level of translation. Our experiments with MG-132, a proteasome inhibitor, show that it is unlikely that the SRP subunits are degraded by the proteasome, suggesting dysregulation of SRP subunits during translation. In recent studies, the concept of coordinated regulation of translation and folding of proteins in oligomers or complexes received strong experimental evidence (Bertolini et al., 2021; Santos et al., 2026; Shiber et al., 2018; Wruck et al., 2025; Wu et al., 2026).

Although the majority of SRP assembly happens in the nucleolus, the heterodimers of the SRP subunits are formed in the cytosol. Thus, they may fold co-translationally together, and the lack of one subunit may dysregulate the translation of the other. Alternatively, they may be degraded by non-proteasome system. It is also possible that they are regulated by a combinatorial mechanism. However, these questions were not in the scope of this work, and they require thorough experimental examination in the future.

During its cycle, SRP recognizes signal sequences at the ribosomes co-translationally and brings the ribosome-polypeptide nascent chain complexes to the SRP receptor in the ER membrane (**Fig. 1**). SRP binds to the ribosomes translating mRNAs of proteins with signal sequences with high affinity of 0.05-0.38 nM (Flanagan et al., 2003). The SRP54 subunit binds signal sequences directly (Kobayashi et al., 2018; Nilsson et al., 2015). Thus, SRP54 plays the major role in the recognition of proteins for transport. Is SRP without SRP54 subunit able to bind the ribosomes? In this work we examined ribosome association with depleted SRP54 through the techniques of crude fractionation and polysome profiling (**Fig. 5**). We show, for the first time, that the loss of SRP54 causes other SRP protein subunits to be shifted to the cytoplasm or pre- monosome fractions, suggesting a loss of SRP binding to the ribosome.

In addition to the protein targeting function, SRP plays a crucial role in protection of mRNAs of secretory and membrane proteins from degradation in the RAPP protein quality control pathway by preventing its activation under normal conditions (Karamyshev and Karamysheva, 2018; Karamyshev et al., 2014; Miller et al., 2026; Pinarbasi et al., 2018; Tikhonova et al., 2022). However, the role of different SRP subunits in this process was not known. In this work, we tested the role of each SRP subunit in mRNA protection for different groups of proteins – secreted proteins into extracellular matrix, proteins with localization in the ER lumen, and plasma membrane proteins (**Fig. 6**). All these proteins are potential substrates for SRP, but they have different final localizations. We have found that the S domain of SRP is responsible for mRNA protection of all above mRNAs regardless of their protein final subcellular localization, while the Alu domain is not, supporting a model of the distinct domain functions. This function is specific to the mRNAs of SRP substrate proteins only; the mRNAs of non-secreted cytosolic proteins were not affected by depletion of any SRP subunit (**Fig. 6**). Our results show that the SRP54 knockdown has the most detrimental effect of all subunits on mRNA of SRP-dependent proteins. Thus, among the S domain subunits, SRP54 has a primary role in mRNA protection, while the role of others decreases with distance from the SRP54 subunit: SRP19 has some effect on mRNA protection, but the loss of the more distant subunits, SRP68/72, was insignificant. Interestingly, the effect of SRP subunit depletion was more noticeable on the mRNAs of extracellular matrix proteins (most sensitive is LGALS3BP mRNA), and slightly less so on ER lumen and plasma membrane proteins. It could be that the latter mRNAs are more stable in general. We also found that protein expression largely correlates with the expression levels of the corresponding mRNAs (**Fig. 6 and 7**). Thus, the S domain of SRP, and especially the SRP54 subunit, plays a major role in protecting secreted and membrane protein mRNAs from abnormal activation of RAPP in the cell.

Our findings about the role of SRP subunits in SRP biogenesis, ribosome-association, and RAPP induction are schematically summarized in **Fig. 8**. Our recent studies show that defects in signal peptide binding to SRP lead to pathological RAPP activation, ultimately causing various human diseases (Gutierrez Guarnizo et al., 2023; Miller et al., 2026; Pinarbasi et al., 2018; Tikhonova et al., 2019). It has also been demonstrated that defects in SRP subunits and their expression are associated with human disease as reviewed in (Kellogg et al., 2022). Furthermore, it has been shown that depletions of SRP19, SRP54, SRP68, and SRP72 cause morphological differences in the *Drosophila* cardiac tissue of (Schroeder et al., 2022), and that SRP may play a role in α-synuclein biogenesis and Parkinson’s disease (Hernandez et al., 2021). Therefore, in addition to a fundamental understanding of SRP biogenesis and the role of its subunits in protein targeting and mRNA protection, our current study provides an important contribution to understanding the molecular mechanisms of human diseases associated with the dysregulation of SRP function.

**Fig. 8.**
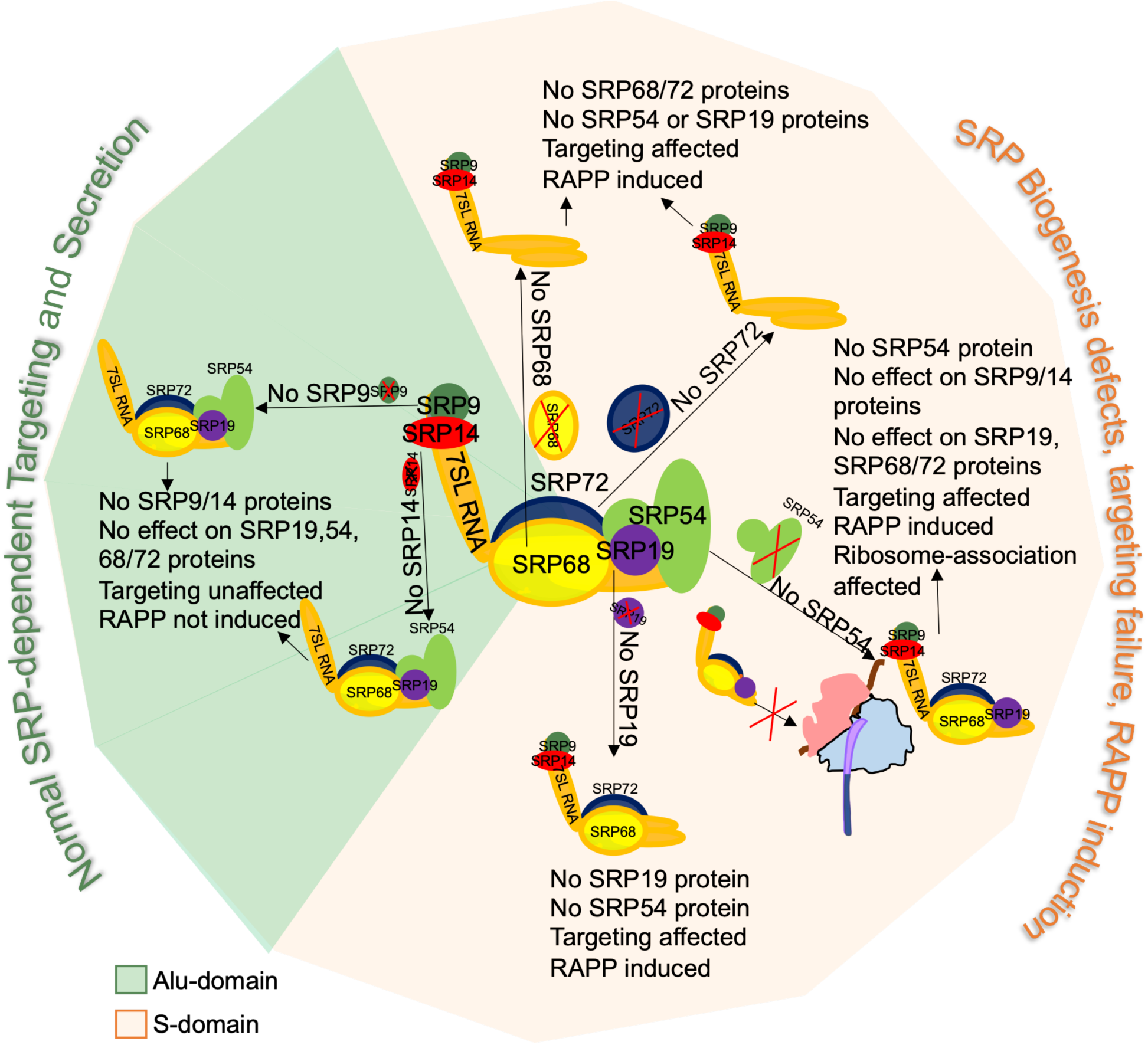
Schematic representation of the effects of SRP subunit defects on its function. When we treated the signal recognition particle with siRNA, we observed a differential impact of subunits on complex formation, interactions with the ribosome, and functional activity. This impact depends on association of the subunits with either the Alu-domain (SRP9/14) or the S- domain (SRP19, SRP54, SRP68/72). Here we present a summary of results emphasizing the effect of SRP subunit depletion. Silencing Alu-domain (light green) SRP9 and SRP14 only lead to SRP9 and SRP14 protein level depletion without any other effects on other SRP protein subunits. Targeting of SRP-dependent proteins is also not affected, nor is RAPP induced. In contrast, we see that there is RAPP induction, SRP-dependent protein targeting failure, and SRP biogenesis defects in the S-domain depletions (light tan). Without SRP68 and SRP72, there is little SRP19 or SRP54 protein expression left. Silencing SRP19 modulates SRP54 protein expression. siSRP19 also leads to SRP-dependent protein targeting failure and RAPP induction. With the loss of SRP54, no other SRP subunit protein levels are depleted, however, there are statistically significant effects to SRP-client protein targeting. Furthermore, the lack of SRP54 leads to the most severe RAPP induction. Finally, silencing SRP54 leads to a failure of the subunits to associate with the ribosome. Therefore, the S-domain of the SRP complex has the most critical effect on SRP biogenesis and function.

## Materials and Methods

### Cell Culture

HeLa Tet-On cells (Clontech) were cultured in Dulbecco’s Modified Eagle’s Medium (DMEM, Sigma) supplemented with 10% Fetal Bovine Serum (FBS) and 1% Penicillin/Streptomycin (P/S, Sigma) at 5% CO2 and 37 °C. Cells were plated at concentrations between 0.6-0.8 x 10^5^ cells/ml approximately 16 hours before transfection with siRNA (Horizon/Dharmacon, **Supplementary Table 1**.) Transfections were performed at a final concentration of 13.75 nM siRNAs with Thermofisher Lipofectamine RNAimax.

### Western Blotting

Media was collected from either 6- or 12-well tissue culture plates and stored at −20°C. Total cell lysate was extracted using a modified Laemmli Sample Buffer supplemented with 1% β-mercaptoethanol and boiled for 5-10 minutes. Proteins were separated into 12% (SRP72, SRP68, SRP54, LGALS3BP, PDIA3, SLC39A14, CHRND, CD248, RPL11, RPS6, and β-Actin), 15% (SRP19, SRP14, SRP9), or 4-15% gradient (K48-linked ubiquitin) SDS-PAGE gels. Proteins were then transferred to 0.45 um PVDF membranes for SRP72, SRP68, SRP54, LGALS3BP, PDIA3, SLC39A14, CHRND, CD248, RPL11, RPS6, and β-Actin, 0.2 µm PVDF membranes for SRP19 and SRP14, or 0.1 µm nitrocellulose membranes for SRP9 using the Biorad Trans-blot Turbo transfer system. SRP9 then underwent fixation on the membrane using 0.4% paraformaldehyde. Blocking and antibody dilutions were performed in 5% dry milk in tris-buffered saline with 0.1% Tween 20 (TBST). All washes were in TBST. Primary and secondary antibodies are described in **Supplementary Table 2**. Relative protein levels were quantified using ImageJ as described (Peterson, 2010). β-actin was used as a loading control and was used for normalization of protein levels in immunoblots using ImageJ.

### Reverse transcriptase quantitative polymerase chain reaction (RT-qPCR)

Total RNA was extracted using TriZol (ThermoFisher) according to manufacturer protocols from 6- or 12-well plates. Experiments were conducted in independent biological replicates as indicated. cDNA was generated using the High Capacity cDNA Reverse Transcription Kit by Applied Biosystems. RT-qPCR was performed on a QuantStudio Real-Time PCR System with SYBR Green Master Mix (ThermoFisher), and relative mRNA levels were calculated by the comparative C_T_ method (or ΔΔC_T_ method) (Schmittgen and Livak, 2008), β-actin was used for normalization. Sequences of primers used are described in **Supplementary Table 3**.

### MG-132Treatment

HeLa Tet-On cells were plated and transfected with siRNA, as described above. Cells were then treated with 10 μM MG-132 or DMSO, incubated for up to 8 hours and collected for analysis by Western blot. Immunoblots were quantified using the ImageJ software described above.

### Crude Fractionation

For crude fractionation, HeLa Tet-On cells were cultured in 10 cm diameter tissue culture plates and treated with 100 μg/ml cycloheximide. HeLa Tet-On cells were then lysed as described for polysome profiling (Karamysheva et al., 2018). Half of the lysate was saved as the input fraction, while the other half was loaded on the 147 mM sucrose cushion before ultracentrifugation at 98000 rpm at +4°C with a TLA100 rotor. The pellet was then resuspended in lysis buffer in the same volume as the supernatant. Equal volumes were loaded for immunoblotting. The fractions (supernatant or pellet) were then analyzed for SRP and ribosomal subunits via Western Blot, as described earlier. The total signal was considered supernatant plus the pellet (S+P). The part was then computed over the whole (S/(S+P) or P/(S+P)) for analysis.

### Polysome Profiling

Polysome profiling was performed as described (Karamysheva et al., 2018). Briefly, cultured HeLa Tet-On cells were cultured in 15 cm diameter tissue culture plates and were treated with 100 μg/ml final concentration of cycloheximide for 15 minutes at 5% CO2 and 37°C. HeLa Tet-On cells were then lysed with the lysis buffer from (Karamysheva et al., 2018). Input Samples were blotted to confirm the absence of SRP54 and the presence of other subunits. HeLa Tet-On cell lysate was centrifuged at 390,000 x g on a 10-50% sucrose gradient, then fractionated and analyzed via the Biocomp Gradient Master and Piston Gradient Fractionator.

Post-processing of fractions included protein precipitation using a 10% final concentration of trichloroacetic acid as described previously (Karamysheva et al., 2018). The total immunoblot signal was considered Free + 40S + 60S + 80S. The part was then computed over the whole (Free/Total, 40S/Total, 60S/Total, or 80S/Total.)

### Statistics

Statistics were performed using the ANOVA, two-way ANOVA, or t-test (as indicated in the figure legend) functions in Excel and GraphPad Prism. Post-hoc tests were performed using GraphPad Prism. Significance was defined as p > 0.05 ns, p<0.05 *, p<0.01 **, p<0.001 ***, p<0.0001 ****. Scatter points indicate independent biological repeats. Error bars represent the standard error of the mean.

## Acknowledgments

This work was supported by the National Institute of General Medical Sciences of the National Institutes of Health under award number R01GM135167 to ALK. The content is solely the authors’ responsibility and does not necessarily represent the official views of the National Institutes of Health.

## References

Aebersold, R., J.N. Agar, I.J. Amster, M.S. Baker, C.R. Bertozzi, E.S. Boja, C.E. Costello, B.F. Cravatt, C. Fenselau, B.A. Garcia, Y. Ge, J. Gunawardena, R.C. Hendrickson, P.J. Hergenrother, C.G. Huber, A.R. Ivanov, O.N. Jensen, M.C. Jewett, N.L. Kelleher, L.L. Kiessling, N.J. Krogan, M.R. Larsen, J.A. Loo, R.R. Ogorzalek Loo, E. Lundberg, M.J. MacCoss, P. Mallick, V.K. Mootha, M. Mrksich, T.W. Muir, S.M. Patrie, J.J. Pesavento, S.J. Petteri, H. Rodriguez, A. Saghatelian, W. Sandoval, H. Schluter, S. Sechi, S.A. Slavoff, L.M. Smith, M.P. Snyder, P.M. Thomas, M. Uhlen, J.E. Van Eyk, M. Vidal, D.R. Walt, F.M. White, E.R. Williams, T. Wohlschlager, V.H. Wysocki, N.A. Yates, N.L. Young, and B. Zhang. 2018. How many human proteoforms are there? Nat Chem Biol. 14:206–214.

Aggeler, R., J. Coons, S.W. Taylor, S.S. Ghosh, J.J. Garcia, R.A. Capaldi, and M.F. Marusich. 2002. A functionally active human F1F0 ATPase can be purified by immunocapture from heart tissue and fibroblast cell lines. Subunit structure and activity studies. J Biol Chem. 277:33906–33912.

Anderson, S., A.T. Bankier, B.G. Barrell, M.H. de Bruijn, A.R. Coulson, J. Drouin, I.C. Eperon, D.P. Nierlich, B.A. Roe, F. Sanger, P.H. Schreier, A.J. Smith, R. Staden, and I.G. Young. 1981. Sequence and organization of the human mitochondrial genome. Nature. 290:457–465.

Becker, M.M., K. Lapouge, B. Segnitz, K. Wild, and I. Sinning. 2017. Structures of human SRP72 complexes provide insights into SRP RNA remodeling and ribosome interaction. Nucleic Acids Res. 45:470–481.

Bellanne-Chantelot, C., B. Schmaltz-Panneau, C. Marty, O. Fenneteau, I. Callebaut, S. Clauin, A. Docet, G.L. Damaj, T. Leblanc, I. Pellier, C. Stoven, S. Souquere, I. Antony-Debr◻, B. Beaupain, N. Aladjidi, V. Barlogis, F. Bauduer, P. Bensaid, O. Boespflug-Tanguy, C. Berger, Y. Bertrand, L. Carausu, C. Fieschi, C. Galambrun, A. Schmidt, H. Journel, F. Mazingue, B. Nelken, T.C. Quah, E. Oksenhendler, M. Ouache, M. Pasquet, V. Saada, F. Suarez, G. Pierron, W. Vainchenker, I. Plo, and J. Donadieu. 2018. Mutations in the SRP54 gene cause severe congenital neutropenia as well as Shwachman-Diamond-like syndrome. Blood. 132:1318–1331.

Bertolini, M., K. Fenzl, I. Kats, F. Wruck, F. Tippmann, J. Schmitt, J.J. Auburger, S. Tans, B. Bukau, and G. Kramer. 2021. Interactions between nascent proteins translated by adjacent ribosomes drive homomer assembly. Science. 371:57–64.

Birse, D.E., U. Kapp, K. Strub, S. Cusack, and A. Aberg. 1997. The crystal structure of the signal recognition particle Alu RNA binding heterodimer, SRP9/14. EMBO J. 16:3757–3766.

Bourdi, M., D. Demady, J.L. Martin, S.K. Jabbour, B.M. Martin, J.W. George, and L.R. Pohl. 1995. cDNA cloning and baculovirus expression of the human liver endoplasmic reticulum P58: characterization as a protein disulfide isomerase isoform, but not as a protease or a carnitine acyltransferase. Arch Biochem Biophys. 323:397–403.

Bousset, L., C. Mary, M.A. Brooks, A. Scherrer, K. Strub, and S. Cusack. 2014. Crystal structure of a signal recognition particle Alu domain in the elongation arrest conformation. RNA. 20:1955–1962.

Brooks, M.A., R.B. Ravelli, A.A. McCarthy, K. Strub, and S. Cusack. 2009. Structure of SRP14 from the Schizosaccharomyces pombe signal recognition particle. Acta Crystallogr D Biol Crystallogr. 65:421–433.

Brown, J.D., B.C. Hann, K.F. Medzihradszky, M. Niwa, A.L. Burlingame, and P. Walter. 1994. Subunits of the Saccharomyces cerevisiae signal recognition particle required for its functional expression. EMBO J. 13:4390–4400.

Carapito, R., M. Konantz, C. Paillard, Z. Miao, A. Pichot, M.S. Leduc, Y. Yang, K.L. Bergstrom, D.H. Mahoney, D.L. Shardy, G. Alsaleh, L. Naegely, A. Kolmer, N. Paul, A. Hanauer, V. Rolli, J.S. Mller, E. Alghisi, L. Sauteur, C. Macquin, A. Morlon, C.S. Sancho, P. Amati-Bonneau, V. Procaccio, A.L. Mosca-Boidron, N. Marle, N. Osmani, O. Lefebvre, J.G. Goetz, S. Unal, N.A. Akarsu, M. Radosavljevic, M.P. Chenard, F. Rialland, A. Grain, M.C. B◻n◻, M. Eveillard, M. Vincent, J. Guy, L. Faivre, C. Thauvin-Robinet, J. Thevenon, K. Myers, M.D. Fleming, A. Shimamura, E. Bottollier- Lemallaz, E. Westhof, C. Lengerke, B. Isidor, and S. Bahram. 2017. Mutations in signal recognition particle SRP54 cause syndromic neutropenia with Shwachman-Diamond-like features. J Clin Invest. 127:4090–4103.

Carden, M.A., J.A. Connelly, E.P. Weinzierl, L.J. Kobrynski, and S. Chandrakasan. 2018. Severe Congenital Neutropenia associated with SRP54 mutation in 22q11.2 Deletion Syndrome: Hematopoietic Stem Cell Transplantation Results in Correction of Neutropenia with Adequate Immune Reconstitution. J Clin Immunol. 38:546–549.

Egea, P.F., J. Napetschnig, P. Walter, and R.M. Stroud. 2008. Structures of SRP54 and SRP19, the two proteins that organize the ribonucleic core of the signal recognition particle from Pyrococcus furiosus. PLoS One. 3:e3528.

Elder, J.T., J. Pan, C.H. Duncan, and S.M. Weissman. 1981. Transcriptional analysis of interspersed repetitive polymerase III transcription units in human DNA. Nucleic Acids Res. 9:1171–1189.

Ercolani, L., B. Florence, M. Denaro, and M. Alexander. 1988. Isolation and complete sequence of a functional human glyceraldehyde-3-phosphate dehydrogenase gene. J Biol Chem. 263:15335–15341.

Flanagan, J.J., J.C. Chen, Y. Miao, Y. Shao, J. Lin, P.E. Bock, and A.E. Johnson. 2003. Signal recognition particle binds to ribosome-bound signal sequences with fluorescence-detected subnanomolar affinity that does not diminish as the nascent chain lengthens. J Biol Chem. 278:18628–18637.

Goldberg, L., A.J. Simon, G. Rechavi, A. Lev, O. Barel, V. Kunik, A. Toren, G. Schiby, H. Tamary, O. Steinberg-Shemer, and R. Somech. 2020. Congenital neutropenia with variable clinical presentation in novel mutation of the SRP54 gene. Pediatr Blood Cancer. 67:e28237.

Grotwinkel, J.T., K. Wild, B. Segnitz, and I. Sinning. 2014. SRP RNA remodeling by SRP68 explains its role in protein translocation. Science. 344:101–104.

Gussakovsky, D., E.P. Booy, M.J.F. Brown, and S.A. McKenna. 2023. Nuclear SRP9/SRP14 heterodimer transcriptionally regulates 7SL and BC200 RNA expression. RNA. 29:1185–1200.

Gutierrez Guarnizo, S.A., M.K. Kellogg, S.C. Miller, E.B. Tikhonova, Z.N. Karamysheva, and A.L. Karamyshev. 2023. Pathogenic signal peptide variants in the human genome. NAR Genom Bioinform. 5:lqad093.

Hainzl, T., S. Huang, and A.E. Sauer-Eriksson. 2002. Structure of the SRP19 RNA complex and implications for signal recognition particle assembly. Nature. 417:767–771.

Hainzl, T., S. Huang, and A.E. Sauer-Eriksson. 2007. Interaction of signal-recognition particle 54 GTPase domain and signal-recognition particle RNA in the free signal-recognition particle. Proc Natl Acad Sci U S A. 104:14911–14916.

Halic, M., T. Becker, M.R. Pool, C.M. Spahn, R.A. Grassucci, J. Frank, and R. Beckmann. 2004. Structure of the signal recognition particle interacting with the elongation-arrested ribosome. Nature. 427:808–814.

Hernandez, S.M., E.B. Tikhonova, K.R. Baca, F. Zhao, X. Zhu, and A.L. Karamyshev. 2021. Unexpected Implication of SRP and AGO2 in Parkinson’s Disease: Involvement in Alpha-Synuclein Biogenesis. Cells. 10.

Iakhiaeva, E., C.S. Hinck, A.P. Hinck, and C. Zwieb. 2009. Characterization of the SRP68/72 interface of human signal recognition particle by systematic site-directed mutagenesis. Protein Sci. 18:2183–2195.

Issa, A., F. Schlotter, J. Flayac, J. Chen, L. Wacheul, M. Philippe, L. Sardini, L. Mostefa, F. Vandermoere, E. Bertrand, C. Verheggen, D.L. Lafontaine, and S. Massenet. 2024. The nucleolar phase of signal recognition particle assembly. Life Sci Alliance. 7.

Juaire, K.D., K. Lapouge, M.M.M. Becker, I. Kotova, M. Michelhans, R. Carapito, K. Wild, S. Bahram, and I. Sinning. 2021. Structural and Functional Impact of SRP54 Mutations Causing Severe Congenital Neutropenia. Structure. 29:15–28 e17.

Karamyshev, A.L., and Z.N. Karamysheva. 2018. Lost in Translation: Ribosome-Associated mRNA and Protein Quality Controls. Front Genet. 9:431.

Karamyshev, A.L., A.E. Patrick, Z.N. Karamysheva, D.S. Griesemer, H. Hudson, S. Tjon-Kon- Sang, I. Nilsson, H. Otto, Q. Liu, S. Rospert, G. von Heijne, A.E. Johnson, and P.J. Thomas. 2014. Inefficient SRP interaction with a nascent chain triggers a mRNA quality control pathway. Cell. 156:146–157.

Karamyshev, A.L., E.B. Tikhonova, and Z.N. Karamysheva. 2020. Translational Control of Secretory Proteins in Health and Disease. Int J Mol Sci. 21.

Karamysheva, Z.N., and A.L. Karamyshev. 2023. Aberrant protein targeting activates quality control on the ribosome. Front Cell Dev Biol. 11:1198184.

Karamysheva, Z.N., E.B. Tikhonova, P.N. Grozdanov, J.C. Huffman, K.R. Baca, A. Karamyshev, R.B. Denison, C.C. MacDonald, K. Zhang, and A.L. Karamyshev. 2018. Polysome Profiling in Leishmania, Human Cells and Mouse Testis. J Vis Exp.

Karamysheva, Z.N., E.B. Tikhonova, and A.L. Karamyshev. 2019. Granulin in Frontotemporal Lobar Degeneration: Molecular Mechanisms of the Disease. Front Neurosci. 13:395.

Kellogg, M.K., S.C. Miller, E.B. Tikhonova, and A.L. Karamyshev. 2021. SRPassing Co-translational Targeting: The Role of the Signal Recognition Particle in Protein Targeting and mRNA Protection. Int J Mol Sci. 22.

Kellogg, M.K., E.B. Tikhonova, and A.L. Karamyshev. 2022. Signal Recognition Particle in Human Diseases. Front Genet. 13:898083.

Kobayashi, K., A. Jomaa, J.H. Lee, S. Chandrasekar, D. Boehringer, S.O. Shan, and N. Ban. 2018. Structure of a prehandover mammalian ribosomal SRP.SRP receptor targeting complex. Science. 360:323–327.

Kramer, A., F. Mulhauser, C. Wersig, K. Groning, and G. Bilbe. 1995. Mammalian splicing factor SF3a120 represents a new member of the SURP family of proteins and is homologous to the essential splicing factor PRP21p of Saccharomyces cerevisiae. RNA. 1:260–272.

Lakkaraju, A.K., C. Mary, A. Scherrer, A.E. Johnson, and K. Strub. 2008. SRP keeps polypeptides translocation-competent by slowing translation to match limiting ER-targeting sites. Cell. 133:440–451.

Linder, M.I., Y. Mizoguchi, S. Hesse, G. Csaba, M. Tatematsu, M. Lyszkiewicz, N. Zietara, T. Jeske, M. Hastreiter, M. Rohlfs, Y. Liu, P. Grabowski, K. Ahomaa, D. Maier-Begandt, M. Schwestka, V. Pazhakh, A.I. Isiaku, B. Briones Miranda, P. Blombery, M.K. Saito, E. Rusha, Z. Alizadeh, Z. Pourpak, M. Kobayashi, N. Rezaei, E. Unal, F. Hauck, M. Drukker, B. Walzog, J. Rappsilber, R. Zimmer, G.J. Lieschke, and C. Klein. 2023. Human genetic defects in SRP19 and SRPRA cause severe congenital neutropenia with distinctive proteome changes. Blood. 141:645–658.

Lutcke, H., S. High, K. Romisch, A.J. Ashford, and B. Dobberstein. 1992. The methionine-rich domain of the 54 kDa subunit of signal recognition particle is sufficient for the interaction with signal sequences. EMBO J. 11:1543–1551.

MacFadyen, J.R., O. Haworth, D. Roberston, D. Hardie, M.T. Webster, H.R. Morris, M. Panico, M. Sutton-Smith, A. Dell, P. van der Geer, D. Wienke, C.D. Buckley, and C.M. Isacke. 2005. Endosialin (TEM1, CD248) is a marker of stromal fibroblasts and is not selectively expressed on tumour endothelium. FEBS Lett. 579:2569–2575.

Majander, A., K. Huoponen, M.L. Savontaus, E. Nikoskelainen, and M. Wikstrom. 1991. Electron transfer properties of NADH:ubiquinone reductase in the ND1/3460 and the ND4/11778 mutations of the Leber hereditary optic neuroretinopathy (LHON). FEBS Lett. 292:289–292.

Mary, C., A. Scherrer, L. Huck, A.K. Lakkaraju, Y. Thomas, A.E. Johnson, and K. Strub. 2010. Residues in SRP9/14 essential for elongation arrest activity of the signal recognition particle define a positively charged functional domain on one side of the protein. RNA. 16:969–979.

Miller, S.C., E.B. Tikhonova, S.M. Hernandez, J.M. Dufour, and A.L. Karamyshev. 2024. Loss of Preproinsulin Interaction with Signal Recognition Particle Activates Protein Quality Control, Decreasing mRNA Stability. J Mol Biol. 436:168492.

Miller, S.C., E.B. Tikhonova, C.C. Rodriguez-Almonacid, C. Bustamante, S.T. Weintraub, and A.L. Karamyshev. 2026. Signal recognition particle-dependent secretome in humans. Sci Rep. 16.

Nilsson, I., P. Lara, T. Hessa, A.E. Johnson, G. von Heijne, and A.L. Karamyshev. 2015. The code for directing proteins for translocation across ER membrane: SRP cotranslationally recognizes specific features of a signal sequence. J Mol Biol. 427:1191–1201.

Oubridge, C., A. Kuglstatter, L. Jovine, and K. Nagai. 2002. Crystal structure of SRP19 in complex with the S domain of SRP RNA and its implication for the assembly of the signal recognition particle. Mol Cell. 9:1251–1261.

Panchal, M., K. Rawat, G. Kumar, K.M. Kibria, S. Singh, M. Kalamuddin, A. Mohmmed, P. Malhotra, and R. Tuteja. 2014. Plasmodium falciparum signal recognition particle components and anti-parasitic effect of ivermectin in blocking nucleo-cytoplasmic shuttling of SRP. Cell Death Dis. 5:e994.

Peterson, T. 2010. Densitometric Analysis using NIH Image. NAVBO eNewsletter. 16.

Petrovic, A., S. Pasqualato, P. Dube, V. Krenn, S. Santaguida, D. Cittaro, S. Monzani, L. Massimiliano, J. Keller, A. Tarricone, A. Maiolica, H. Stark, and A. Musacchio. 2010. The MIS12 complex is a protein interaction hub for outer kinetochore assembly. J Cell Biol. 190:835–852.

Piazzon, N., F. Schlotter, S. Lefebvre, M. Dodre, A. Mereau, J. Soret, A. Besse, M. Barkats, R. Bordonne, C. Branlant, and S. Massenet. 2013. Implication of the SMN complex in the biogenesis and steady state level of the signal recognition particle. Nucleic Acids Res. 41:1255–1272.

Pinarbasi, E.S., A.L. Karamyshev, E.B. Tikhonova, I.H. Wu, H. Hudson, and P.J. Thomas. 2018. Pathogenic Signal Sequence Mutations in Progranulin Disrupt SRP Interactions Required for mRNA Stability. Cell Rep. 23:2844–2851.

Pohl, C., and I. Dikic. 2019. Cellular quality control by the ubiquitin-proteasome system and autophagy. Science. 366:818–822.

Poritz, M.A., H.D. Bernstein, K. Strub, D. Zopf, H. Wilhelm, and P. Walter. 1990. An E. coli ribonucleoprotein containing 4.5S RNA resembles mammalian signal recognition particle. Science. 250:1111–1117.

Ren, Y.G., K.W. Wagner, D.A. Knee, P. Aza-Blanc, M. Nasoff, and Q.L. Deveraux. 2004. Differential regulation of the TRAIL death receptors DR4 and DR5 by the signal recognition particle. Mol Biol Cell. 15:5064–5074.

Rieser, E., S.M. Cordier, and H. Walczak. 2013. Linear ubiquitination: a newly discovered regulator of cell signalling. Trends Biochem Sci. 38:94–102.

Romisch, K., J. Webb, J. Herz, S. Prehn, R. Frank, M. Vingron, and B. Dobberstein. 1989. Homology of 54K protein of signal-recognition particle, docking protein and two E. coli proteins with putative GTP-binding domains. Nature. 340:478–482.

Safina, A., H. Garcia, M. Commane, O. Guryanova, S. Degan, K. Kolesnikova, and K.V. Gurova. 2013. Complex mutual regulation of facilitates chromatin transcription (FACT) subunits on both mRNA and protein levels in human cells. Cell Cycle. 12:2423–2434.

Santos, J., S.J. Tans, N. Ban, G. Kramer, and B. Bukau. 2026. Cotranslational Assembly of Oligomeric Proteins. Annu Rev Biochem.

Schmaltz-Panneau, B., A. Pagnier, S. Clauin, J. Buratti, C. Marty, O. Fenneteau, K. Dieterich, B. Beaupain, J. Donadieu, I. Plo, and C. Bellanne-Chantelot. 2021. Identification of biallelic germline variants of SRP68 in a sporadic case with severe congenital neutropenia. Haematologica. 106:1216–1219.

Schmittgen, T.D., and K.J. Livak. 2008. Analyzing real-time PCR data by the comparative C(T) method. Nat Protoc. 3:1101–1108.

Schroeder, A.M., T. Nielsen, M. Lynott, G. Vogler, A.R. Colas, and R. Bodmer. 2022. Nascent polypeptide-Associated Complex and Signal Recognition Particle have cardiac-specific roles in heart development and remodeling. PLoS Genet. 18:e1010448.

Schroeder, B.C., S. Waldegger, S. Fehr, M. Bleich, R. Warth, R. Greger, and T.J. Jentsch. 2000. A constitutively open potassium channel formed by KCNQ1 and KCNE3. Nature. 403:196–199.

Schurch, C., T. Schaefer, J.S. Muller, P. Hanns, M. Arnone, A. Dumlin, J. Scharer, I. Sinning, K. Wild, J. Skokowa, K. Welte, R. Carapito, S. Bahram, M. Konantz, and C. Lengerke. 2021. SRP54 mutations induce congenital neutropenia via dominant-negative effects on XBP1 splicing. Blood. 137:1340–1352.

Schwanhausser, B., D. Busse, N. Li, G. Dittmar, J. Schuchhardt, J. Wolf, W. Chen, and M. Selbach. 2011. Global quantification of mammalian gene expression control. Nature. 473:337–342.

Seegmiller, J.E., F.M. Rosenbloom, and W.N. Kelley. 1967. Enzyme defect associated with a sex-linked human neurological disorder and excessive purine synthesis. Science. 155:1682–1684.

Shiber, A., K. Doring, U. Friedrich, K. Klann, D. Merker, M. Zedan, F. Tippmann, G. Kramer, and B. Bukau. 2018. Cotranslational assembly of protein complexes in eukaryotes revealed by ribosome profiling. Nature. 561:268–272.

Siegel, V., and P. Walter. 1986. Removal of the Alu structural domain from signal recognition particle leaves its protein translocation activity intact. Nature. 320:81–84.

Siegel, V., and P. Walter. 1988. Each of the activities of signal recognition particle (SRP) is contained within a distinct domain: analysis of biochemical mutants of SRP. Cell. 52:39–49.

Soni, K., G. Kempf, K. Manalastas-Cantos, A. Hendricks, D. Flemming, J. Guizetti, B. Simon, F. Frischknecht, D.I. Svergun, K. Wild, and I. Sinning. 2021. Structural analysis of the SRP Alu domain from Plasmodium falciparum reveals a non-canonical open conformation. Commun Biol. 4:600.

Taylor, K.M., H.E. Morgan, A. Johnson, and R.I. Nicholson. 2005. Structure-function analysis of a novel member of the LIV-1 subfamily of zinc transporters, ZIP14. *FEBS Lett*. 579:427- 432.

Tikhonova, E.B., S.A. Gutierrez Guarnizo, M.K. Kellogg, A. Karamyshev, I.M. Dozmorov, Z.N. Karamysheva, and A.L. Karamyshev. 2022. Defective Human SRP Induces Protein Quality Control and Triggers Stress Response. J Mol Biol. 434:167832.

Tikhonova, E.B., Z.N. Karamysheva, G. von Heijne, and A.L. Karamyshev. 2019. Silencing of Aberrant Secretory Protein Expression by Disease-Associated Mutations. J Mol Biol. 431:2567–2580.

Tinari, N., I. Kuwabara, M.E. Huflejt, P.F. Shen, S. Iacobelli, and F.T. Liu. 2001. Glycoprotein 90K/MAC-2BP interacts with galectin-1 and mediates galectin-1-induced cell aggregation. Int J Cancer. 91:167–172.

Tuteja, R. 2007. Unraveling the components of protein translocation pathway in human malaria parasite Plasmodium falciparum. Arch Biochem Biophys. 467:249–260.

Uhlen, M., L. Fagerberg, B.M. Hallstrom, C. Lindskog, P. Oksvold, A. Mardinoglu, A. Sivertsson, C. Kampf, E. Sjostedt, A. Asplund, I. Olsson, K. Edlund, E. Lundberg, S. Navani, C.A. Szigyarto, J. Odeberg, D. Djureinovic, J.O. Takanen, S. Hober, T. Alm, P.H. Edqvist, H. Berling, H. Tegel, J. Mulder, J. Rockberg, P. Nilsson, J.M. Schwenk, M. Hamsten, K. von Feilitzen, M. Forsberg, L. Persson, F. Johansson, M. Zwahlen, G. von Heijne, J. Nielsen, and F. Ponten. 2015. Proteomics. Tissue-based map of the human proteome. Science. 347:1260419.

Uhlen, M., M.J. Karlsson, A. Hober, A.S. Svensson, J. Scheffel, D. Kotol, W. Zhong, A. Tebani, L. Strandberg, F. Edfors, E. Sjostedt, J. Mulder, A. Mardinoglu, A. Berling, S. Ekblad, M. Dannemeyer, S. Kanje, J. Rockberg, M. Lundqvist, M. Malm, A.L. Volk, P. Nilsson, A. Manberg, T. Dodig-Crnkovic, E. Pin, M. Zwahlen, P. Oksvold, K. von Feilitzen, R.S. Haussler, M.G. Hong, C. Lindskog, F. Ponten, B. Katona, J. Vuu, E. Lindstrom, J. Nielsen, J. Robinson, B. Ayoglu, D. Mahdessian, D. Sullivan, P. Thul, F. Danielsson, C. Stadler, E. Lundberg, G. Bergstrom, A. Gummesson, B.G. Voldborg, H. Tegel, S. Hober, B. Forsstrom, J.M. Schwenk, L. Fagerberg, and A. Sivertsson. 2019. The human secretome. Sci Signal. 12. Volz, A., E. Weiss, J. Trowsdale, and A. Ziegler. 1994. Presence of an expressed beta-tubulin gene (TUBB) in the HLA class I region may provide the genetic basis for HLA-linked microtubule dysfunction. Hum Genet. 93:42–46.

Weichenrieder, O., K. Wild, K. Strub, and S. Cusack. 2000. Structure and assembly of the Alu domain of the mammalian signal recognition particle. Nature. 408:167–173.

Wild, K., G. Bange, D. Motiejunas, J. Kribelbauer, A. Hendricks, B. Segnitz, R.C. Wade, and I. Sinning. 2016. Structural Basis for Conserved Regulation and Adaptation of the Signal Recognition Particle Targeting Complex. J Mol Biol. 428:2880–2897.

Wild, K., K.D. Juaire, K. Soni, V. Shanmuganathan, A. Hendricks, B. Segnitz, R. Beckmann, and I. Sinning. 2019. Reconstitution of the human SRP system and quantitative and systematic analysis of its ribosome interactions. Nucleic Acids Res. 47:3184–3196.

Wruck, F., J. Schmitt, K. Till, K. Fenzl, M. Bertolini, F. Tippmann, A. Katranidis, B. Bukau, G. Kramer, and S.J. Tans. 2025. Co-translational ribosome pairing enables native assembly of misfolding-prone subunits. Nat Commun. 16:7626.

Wu, X., H. Zhang, L. Huang, S. Zhang, J. He, S. Wang, W. Zhu, Y. Li, J. Zhou, and X.M. Liu. 2026. Cotranslational assembly directs the biogenesis of the m(6)A methyltransferase complex. Proc Natl Acad Sci U S A. 123:e2517258123.

