## Supplementary Tables for "Human SRP Subunits Coordinate Protein Targeting and mRNA Quality Control"

**Supplementary Table 1. siRNA used in this study**

| siRNA | Company | Catalog Number/Sequence | Reference |
|---|---|---|---|
| siSRP9 | Horizon/Dharmacon | L-019731-01-0005 | This study |
| siSRP14 | Horizon/Dharmacon | L-017767-01-0005 | This study |
| siSRP19 | Horizon/Dharmacon | L-019729-01-0005 | This study |
| siSRP54 | Horizon/Dharmacon | 5'GAAAUGAACAGGAGUCAAUdTdT3' | (Ren et al., 2004) |
| siSRP68 | Horizon/Dharmacon | L-005121-02-0005 | This study |
| siSRP72 | Horizon/Dharmacon | L-019716-00-0005 | This study |

**Supplementary Table 2. Antibodies used in this study**

| Antibody | Company | Catalog Number |
|---|---|---|
| Anti-SRP9 | Novus Biologicals | NBP2-61107 |
| Anti-SRP14 | Proteintech | 11528-1-AP |
| Anti-SRP19 | ABclonal | A6752 |
| Anti-SRP54 | BD Bioscience | 610941 |
| Anti-SRP68 | Proteintech | 11585-1-AP |
| Anti-SRP72 | Abcam | Ab200199 |
| Anti-RPS6 | Santa Cruz | Sc-74459 |
| Anti-RPL11 | Abcam | Ab79352 |
| Goat anti-mouse | Jackson Laboratories | 115-035-003 |
| Goat anti-rabbit | Cytiva | NA934 |
| Anti-β-actin | Proteintech | 66009-1-Ig |
| Anti-PDIA3 | Proteintech | 15967-1-AP |
| Anti-SLC39A14 (ZIP14) | ABclonal | A10413 |
| Anti-LGALS3BP | Proteintech | 10281-1-AP |
| Anti-CHRND | ABclonal | A10467 |
| Anti-CD248 | Proteintech | 60170-1-Ig |

**Supplementary Table 3. RT qPCR Primers used in this study**

| Gene Name | Forward Primer | Reverse Primer | Reference |
|---|---|---|---|
| SRP9 | CAGATGATTTAGTTTGTTTGGTGT | GTAACATTGCGGGCTTCCTT | This study |
| SRP14 | ATGGTGTTGTTGGAGAGCGA | AGGTGATATAGACGCTGCCC | This study |
| SRP19 | GCCGACCAGGACAGGTTTAT | CTTCCCTCTGCGATGGTCTT | This study |
| SRP54 | ACACCCGATCCTCTTGCTAC | CAGTACGGATGGTGCCAAAG | (Karamyshev et al., 2014) |
| SRP68 | CCAGTCAGCCAAACAGGCA | GACACTCGCTGAGCATTGAT | This study |
| SRP72 | GCTGAACCAGGCCATGAAAA | GACATTTTGGTCAGTGTCTTCTG | This study |
| LGALS3BP | CCTGTCGTCAGTCAAGTGCT | GGAGGATGGCAAAGAGGCTT | This study |
| LOX | CCAGCAGATCCAATGGGAGAA | CTGAGGCTGGTACTGTGAGC | (Tikhonova et al., 2022) |
| MMP2 | AGTCTGTGTTGTCCAGAGGC | TGAAGCCAAGCGGTCTAAGT | This study |
| SPOCK1 | GGACAAGTACTGGAACCGC | ACACACACTTTGTGAGGGCT | (Tikhonova et al., 2022) |
| CALR | CAGTTCCGGCAAGTTCTACG | GAAACTGGCCGACAGAGCAT | (Karamyshev et al., 2014) |
| HSPA5 | TGCTATTGCTTATGGCCTGGA | GAGACACATCGAAGGTTCCG | (Karamyshev et al., 2014) |
| HYOU1 | GGAGAAGCAGGAACGGGAAA | GAGATCTCCTCACGCTGCTC | (Tikhonova et al., 2022) |
| PDIA3 | GATGGGCCTGTGAAGGTAGT | GGGCTCCAGGTTCTTACAGT | (Tikhonova et al., 2022) |
| ALPI | GAAGGTCGCCAAGAACCTCAT | TTCTTCTGCCCCTTTAGGATC | (Karamyshev et al., 2014) |
| CD24 | CTCCTACCCACGCAGATTTATTC | AGAGTGAGACCACGAAGAGAC | (Tikhonova et al., 2022) |
| CD248 | CCAGACCACCACTCATTTGC | TCTGAGGACAAGGGCATCTG | (Tikhonova et al., 2022) |
| CHRND | ATGTGTGGATAGAGCACGGC | GCGCAGGACACTGATGTTTC | (Tikhonova et al., 2022) |
| HLAB | CAGTTCGTGAGGTTCGACAG | CAGCCGTACATGCTCTGGA | This study |
| KCNQ1 | GCCACCTCAACCTCATGGTG | GCGATCCTTGCTCTTTTCTGAG | (Tikhonova et al., 2022) |
| SLC39A14 | TCCGAGCGCCAGGTTTATTC | CCATAAGCCAAGCAGGGTCA | This study |
| HPRT | CTGAGGATTTGGAAAGGGTGTT | ATCTCCTTCATCACATCTCGAG | (Karamyshev et al., 2014) |
| SRPRA | TGCCGCCATGCTCGACTTCT | CTGCAGCAGCACGGAACGAA | (Karamyshev et al., 2014) |
| GAPDH | GTCTCCTCTGACTTCAACAGCG | ACCACCCTGTTGCTGTAGCCAA | (Karamyshev et al., 2014) |
| ACTB | CACCTTCTACAATGAGCTGCG | TAGCACAGCCTGGATAGCAAC | (Karamyshev et al., 2014) |

### Supplementary Table 4. Housekeeping Genes and Protein Function

| Gene | Common Name | Function | Source |
|---|---|---|---|
| SRP-independent or non-SRP targeted clients with cytoplasm localization | | | |
| ACTB | β-actin | Cytoskeleton | |
| HPRT | Hypoxanthine-guanine phosphoribosyltransferase | Purine salvage pathway | (Seegmiller et al., 1967) |
| GAPDH | Glyceraldehyde-3-phosphate dehydrogenase | Glycolysis | (Ercolani et al., 1988) |
| NSL1 | NSL1 component of MIS12 kinetochore complex | Mitosis | (Petrovic et al., 2010) |
| TUBB | β-tubulin | Cytoskeleton | (Volz et al., 1994) |
| Non-SRP targeted clients with mitochondrial localization | | | |
| MTCO1 | Cytochrome c oxidase subunit 1 | Oxidative phosphorylation | (Anderson et al., 1981) |
| MTND1 | NADH-ubiquinone oxidoreductase chain 1 | Complex I biogenesis; electron transport chain | (Majander et al., 1991) |
| MTATP8 | ATP synthase protein 8 | Hydrogen ion transport; electron transport chain | (Aggeler et al., 2002) |
| Non-SRP targeted clients with nuclear localization | | | |
| SF3A1 | Splicing Factor 3a subunit 1 | Spliceosome assembly and pre-mRNA splicing | (Kramer et al., 1995) |

### Supplementary Table 5. Genes and Protein Function for SRP-targeted Clients

| Gene | Common Name | Function | Source |
|---|---|---|---|
| Extracellular matrix localization | | | |
| LGALS3BP | galectin-3-binding protein | Cell-cell adhesion and immune response | (Tinari et al., 2001) |
| Endoplasmic reticulum localization | | | |
| PDIA3 | Protein disulfide isomerase A3 | Catalyzes disulfide bonds, ER chaperone | (Bourdi et al., 1995) |
| Plasma membrane localization | | | |
| CD248 | Endosialin, TEM1 | Angiogenesis | (MacFadyen et al., 2005) |
| KCNQ1 | Potassium voltage-gated channel subfamily KQT member 1 | Voltage-gated potassium ion channel | (Schroeder et al., 2000) |
| SLC39A14 | Metal Cation symporter ZIP14 | Divalent zinc ion channel | (Taylor et al., 2005) |
